# Enhanced Production of HCV E1E2 Subunit Vaccine Candidates via Protein-Protein Interaction Identification in Glycoengineered CHO cells

**DOI:** 10.1101/2025.05.20.655202

**Authors:** Mina Ying Min Wu, Frances Rocamora, Caressa M. Robinson, Seunghyeon Shin, Eric A. Toth, Thomas R. Fuerst, Svetlana Maurya, Nathan E. Lewis

## Abstract

Hepatitis C Virus (HCV) is a bloodborne virus that affects 57 million people globally with infections that can often go unnoticed, and it is the leading cause of chronic liver disease and cancer. Thus, development of an HCV vaccine is a major medical and public health concern. While prior work has developed secreted E1E2 (sE1E2) protein vaccine candidates, efforts to express it recombinantly in Chinese hamster ovary (CHO) cells have resulted in very low titers. To address this challenge, here we employed a multi-omics approach to identify protein interactors that enhance the secretion of sE1E2. By detecting Protein-Protein Interactions (PPIs) using Biotinylation by Antibody Recognition (BAR) and integrating the data with RNA-Seq, we identified proteins within the secretory pathway that interact with sE1E2 and validated their impact by overexpressing the interacting proteins. Among these, CUL4A and YWHAH enhanced sE1E2 secretion in glycoengineered CHO (geCHO) cells. The integration of omics techniques and genetic engineering in this study provides valuable insights into improving protein secretion in CHO cells, paving the way for the development of more affordable and accessible biotherapeutics.

## Introduction

Hepatitis C Virus (HCV) is the leading cause of chronic liver disease such as cirrhosis and hepatocellular carcinoma (Khullar & Firpi, 2015), making the development of a vaccine critically important. Although HCV can be treated with direct-acting antivirals (DAAs) such as sofosbuvir/velpatasvir over the course of a few months, the viral damage is often already done. Moreover, DAAs do not prevent reinfections or transmissions, since infections often go unnoticed (Manns et al., 2017). An effective vaccine for HCV has yet to be achieved due to several factors. First, the error-prone RNA replication process of HCV leads to high mutation rates, resulting in at least 7 genotypes and over 90 subtypes of HCV (Duncan et al., 2020; Guest & Pierce, 2021). Second, viral glycans shield important neutralizing epitopes on the E1E2 heterodimer complex surface (Hedskog et al., 2019; Peña et al., 2022). Third, immunodominant, non-neutralizing epitopes distract the immune response from generating broadly neutralizing antibodies (bNAbs) (Toth et al., 2021). Fourth, it remains challenging to produce a homogeneous E1E2 vaccine at scale, given the difficulty of generating large quantities of subunit vaccine protein (Toth et al., 2021; Lavie et al., 2006). Therefore, there remain multiple challenges to vaccine development, but it is critical since a vaccine could provide a preventative measure for millions of people.

Recent advances in new technology platforms exist that can be applied to develop more efficacious vaccines. For example, changes in glycosylation may enhance vaccine efficacy (Deng et al., 2022; Liu et al., 2016). Glycosylation can impact protein stability and half-life, which in turn can enhance efficacy and immune responses (Rocamora et al., 2023). In addition, removing glycosylation sites on the protein can expose important epitope sites for neutralizing antibody (NAb) binding. Also, by adding glycans to cover up known neutralization binding sites while simultaneously removing other specific N-glycan sites, new binding regions can be exposed (Ringe et al., 2019). Thus, tuning glycosylation on vaccine proteins with geCHO cells could help elicit Nabs (Kulakova et al., 2025).

Consistent with the evidence that glycosylation impacts vaccine efficacy, we previously found that glycoengineered CHO (geCHO) cells can enhance neutralization of a secreted E2 (sE2) vaccine candidate (Kulakova et al., 2025), particularly one glycoform we called geCHO.sE2.1, produced in a geCHO cell line "Cell Line 1" (CL1). CL1 was genetically modified to produce simple glycans that are monoantennary, asialylated and fucosylated on glycoproteins with glycosyltransferase knockouts (KO) for *Mgat2, St3gal3/4/6, B3gnt2,* and *Sppl3*. In contrast, our control (CTRL) CHO cells produce branched and complex glycans, as they only harbor a *Sppl3* KO.

Aside from glycoengineering, other protein engineering strategies can further enhance vaccine design. For example, an improved HCV vaccine candidate was developed by including the E1 subunit alongside E2, forming an E1E2 heterodimer vaccine that mimics the glycoprotein on the surface of HCV (Guest et al., 2021; Wang et al., 2022; Metcalf et al., 2023). The native membrane-bound E1E2 (mbE1E2) heterodimer on the virus surface is the target for vaccine designs to induce bNAbs and stimulate B cell immunity (Lavie et al., 2007; Peña et al., 2022; Toth et al., 2021). The E1E2 complex vaccine presents conserved antigenic epitope sites on E2 including domains A, B/AR3, D, and E; and antigenic epitope sites on E1 including the N-terminal residues 192-202 and a conserved alpha-helix 314-327 (Toth et al., 2023). In addition, the E1E2 heterodimer structure presents additional antigenic regions such as AR4 and AR5 that require intact E1E2 for effective antibody recognition (Pierce et al., 2024). Studies in chimpanzees have shown that vaccination with the E1E2 complex elicits a stronger immune response compared to E2 alone (Frey et al., 2010; Kundu et al., 2024). Additionally, AR4A is associated with the clearance of the virus (Frumento et al., 2022; Kinchen et al., 2019).

With E1E2 being a target for a protein-based vaccine, there is a clear need for its mass production in high quantities, especially since the native E1E2 heterodimer is a membrane-bound protein. Current production of membrane-bound E1E2 (mbE1E2) in CHO cells yields only 1 mg of purified mbE1E2 per 100g of CHO cells (Logan, et al. 2016) and 2-5 mg per liter in suspension HEK cells (Toth, et al. 2021). In addition, non-uniformity of mbE1E2 can complicate downstream processing and make quality control difficult (Guest et al., 2021). Fortunately, secreted versions of E1E2 have been created by replacing the transmembrane domain (TMD) with natural and synthetic scaffolds such as leucine zipper and SynZip (SZ) (Guest, et al., 2021; Metcalf et al., 2023). Therefore, a secreted version of E1E2 (sE1E2.SZ) was created with a furin cleavage site containing six arginines between the E1 and E2 ectodomains. This facilitates proper cleavage and encourages native-like assembly (Metcalf et al., 2023). Additionally, a stabilizing mutation (H445P) in Domain D was incorporated into the sE1E2 design expressed in this study, sE1E2.SZ-H445P. This mutation is located in a region lacking secondary structure near key antibody-binding sites and has been shown to improve antibody binding stability by reducing the epitope site’s mobility (Pierce et al., 2020).

With the promising results of an E1E2-based vaccine, we expressed the sE1E2.SZ-H445P in geCHO cells, specifically in CTL and CL1 cell lines. However, this resulted in insufficient amounts of protein to purify and use for binding affinity and antigenicity tests. To improve secretion of the E1E2 protein, we measured its Protein-Protein Interactions (PPIs) using an innovative method, biotinylation by antibody recognition (BAR) (Bar et al., 2018; Masson et al., 2024). In the BAR method here, sE1E2 cells were fixed and permeabilized, and then incubated with anti-E1E2 antibodies. Then a secondary antibody, conjugated with horseradish peroxidase, was applied, along with H_2_O_2_ and tyramide-biotin. This biotinylated the proteins interacting with sE1E2. Biotinylated proteins were then pulled down and identified via mass spectrometry (MS). We applied BAR and RNA-Seq to the geCHO cells expressing E1E2 to identify protein interactors, especially in the secretory pathway, such as those in the ER and Golgi apparatus involved in translocation, protein folding, vesicle transport, quality control, and post-translational modifications (PTMs). We overexpressed a selection of significant PPI, and found two proteins, CUL4A and YWHAH, significantly improved sE1E2 secretion when overexpressed in geCHO cells. Thus, using PPI and transcriptomic data, we identified PPIs that improve the production of this difficult-to-express protein.

## Results

### Generating stable cell clones expressing sE1E2.SZ-H445P

Plasmids harboring genes for furin protease and sE1E2.SZ-H445P were sequentially transfected into the control (CTRL) and CL1 (Figure 1A) cell lines using Neon electroporation transfection to generate stable pools expressing sE1E2. After transfection, cloning was performed for each cell line, and individual clones were screened for sE1E2 secretion via dot blot analysis. Two clones from each cell line were selected for PPI analysis and transcriptomic studies: CTRL(H445P:Furin) #1, CTRL(H445P:Furin) #2, CL1(H445P:Furin) #17, and CL1(H445P:Furin) #21.

**Figure 1.**
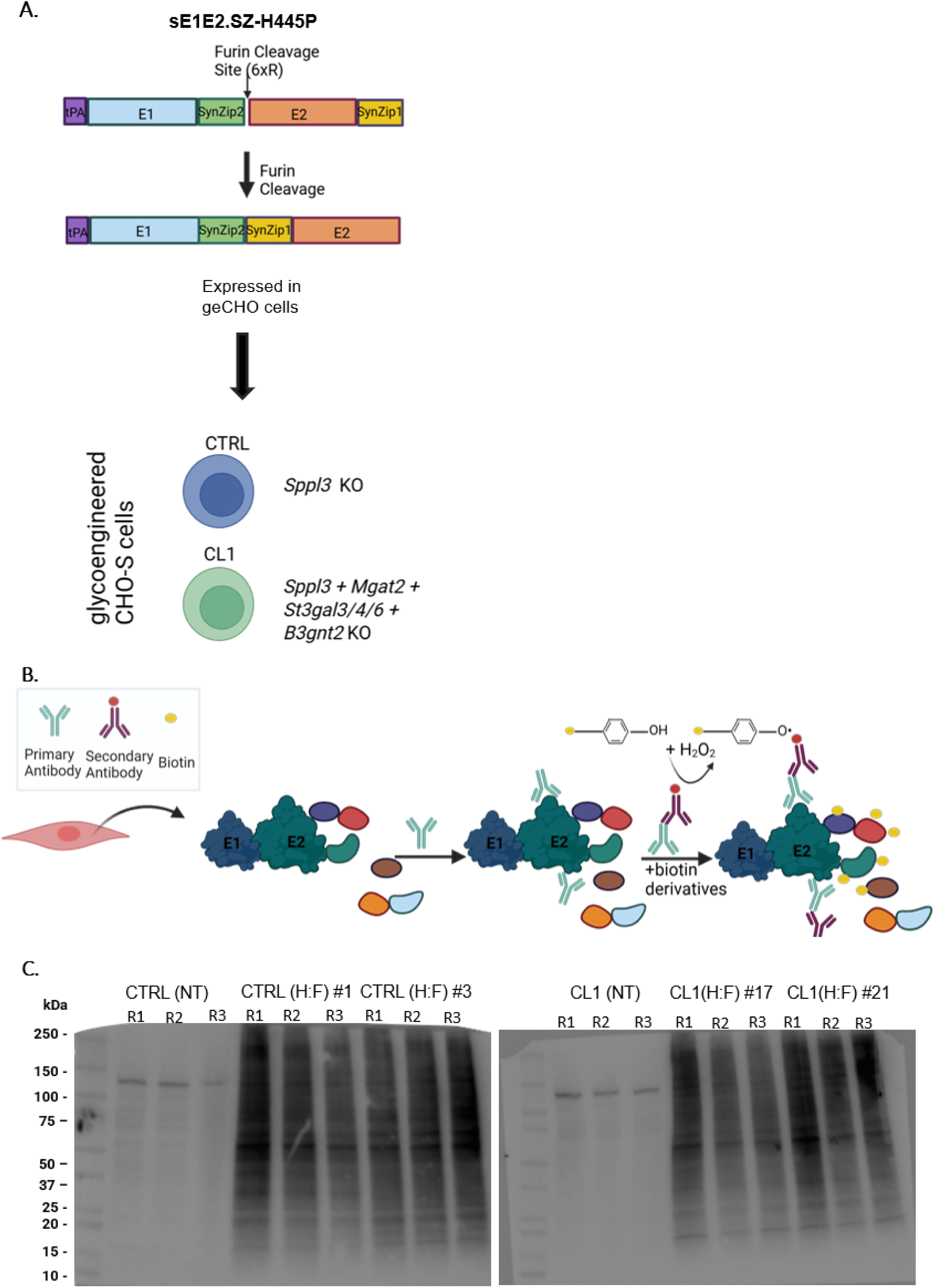
Schematic of the sE1E2.sz-H445p plasmid design and biotinylation confirmation in BAR samples. **(A)** The sE1E2.sz-H445p plasmid was designed for secretion by replacing the transmembrane domains (TMD) with a synthetic scaffold (SynZip), which lacks human sequence homology. Furin co-expression was incorporated to cleave the 6xArginine sites, facilitating native-like assembly. The construct includes a key proline substitution at position H445 (H445p) in Domain D of E2. This plasmid was co-expressed with human furin in two geCHO cell lines: CTRL (*Sppl3* knockout) and CL1 (*Sppl3, Mgat2, St3gal3/4/6, B3gnt2* knockout). **(B)** Stable clones of wild-type and CL1 were obtained expressing sE1E2.sz-H445P. Two clones from each geCHO cell line were selected for biotinylation by antibody recognition (BAR) analysis. Cells were fixed, permeabilized, and stained with the primary HCV1 antibody targeting Domain D of E2. A secondary anti-human horseradish peroxidase (HRP)-conjugated antibody was used to bind HCV1. Upon addition of biotin and hydrogen peroxide, proteins proximal to E1E2 were biotinylated. **(C)** BAR was performed on technical triplicates of each clone, with biotin added for 5 minutes. Biotinylation was confirmed by western blotting, with 25 µg of lysates loaded per sample. Streptavidin-HRP staining was used to visualize biotinylated proteins. Non-transfected parental cell lines served as negative controls.

A Western blot was performed under reducing conditions for each clone and probed with an anti-E1 antibody to confirm secretion of sE1E2. Supernatant samples were collected daily over four days, and cell lysate was collected on day 4 (Supplementary Figure S1A). Both cleaved forms of sE1E2 (processed by furin) and uncleaved forms of sE1E2 (not processed by furin) were observed in the supernatant and the lysate. The cleaved sE1E2 appeared as either E1 (∼30 kDa) or E2 (∼65 kDa), depending on the antibody used for detection. In contrast, the uncleaved sE1E2 was detected at ∼100 kDa.

We attempted various methods to improve furin-mediated cleavage of sE1E2, including codon optimization of furin specifically for CHO cells and placing both furin and sE1E2 genes on the same plasmid construct. Despite these attempts, uncleaved forms of sE1E2 persisted, suggesting either a furin functionality issue or inaccessibility of the furin cleavage site for efficient proteolytic processing.

### BAR targeting proximate proteins of E1E2 produced in geCHO cells

We used BAR to study PPIs in stable clones expressing sE1E2.SZ-H445P. Unlike other techniques like BioID (Kim & Roux, 2016; Pfeiffer et al., 2022; Sears et al., 2019; Samoudi et al., 2021) — which requires fusion of the protein of interest with a BirA enzyme. It also requires lengthy labeling periods that may compromise protein functions — BAR preserves cell integrity, requiring only fixation and treatment with readily available antibodies. BAR employs an HRP conjugated antibody specific to the target protein to biotinylate nearby proteins, capturing proximal proteins at a specific moment during harvesting.

To perform BAR on the clones, cells were fixed, permeabilized, and labeled with HCV1 primary antibody targeting domain E of the E2 protein (Figure 1B). We then applied a secondary anti-human HRP-conjugated antibody to bind to HCV1. Upon biotin addition, HRP transfers biotin to proteins in proximity to sE1E2. Western blot analysis with HRP-streptavidin confirmed successful biotinylation, showing significantly more labeled proteins in sE1E2-expressing clones compared to non-transfected controls (Figure 1C). We then isolated biotinylated samples using streptavidin-coated beads, performed trypsin digestion, and analyzed the results by liquid chromatography-tandem mass spectrometry (LC-MS/MS).

### Preprocessing and quality control analysis of proximate proteins of E1E2 in the BAR proteomic data

The BAR proteomics spectra were analyzed using MaxQuant software (Tyanova, Temu, & Cox, 2016), and the MS/MS spectra were searched against the *Cricetulus griseus* Uniprot protein sequence database. Proteomic data was analyzed using the DEP package (Zhang et al., 2018) in R Studio. Our pre-processing analysis of plotting the total proteins against each sample and its replicates showed consistent protein detection across samples with the nonproducer cell lines containing less protein, as expected (Figure 2A). We normalized the sample using variance stabilizing transformation (Zhang et al., 2018), which stabilizes the variance across the entire range of expression levels. A principal component analysis (PCA) revealed clear separation between producers and nonproducers — principal component 1 accounted for 70% of the variance (Figure 2B). Following quality control, the missing values were imputed using left-shifted Gaussian Distribution for imputation with missing not at random (MNAR) (Zhang et al., 2018), which shifts the imputed values toward the lower end of the detection range. It imputes values based on the assumption that missing values are likely to be low-abundance proteins.

**Figure 2:**
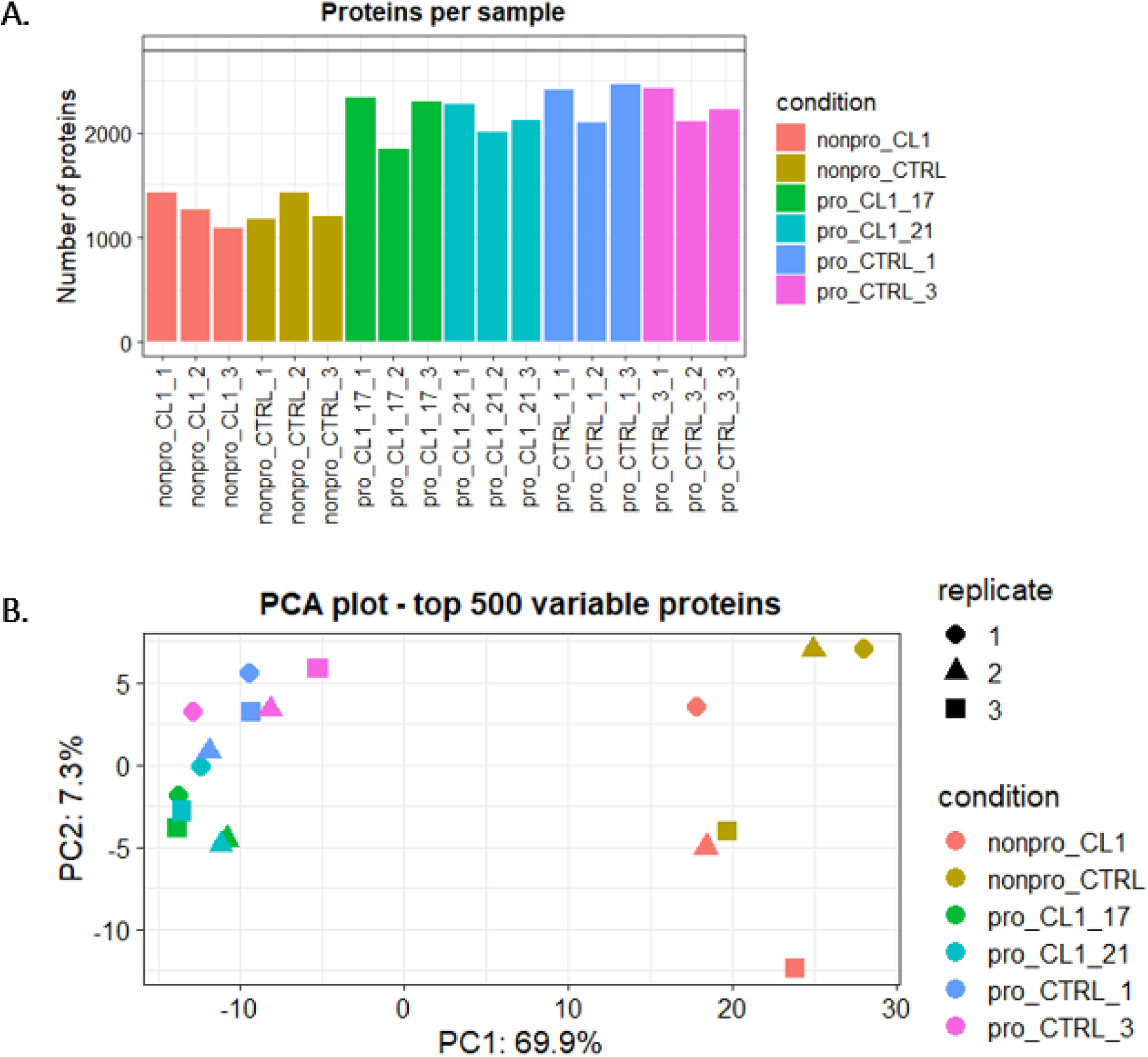
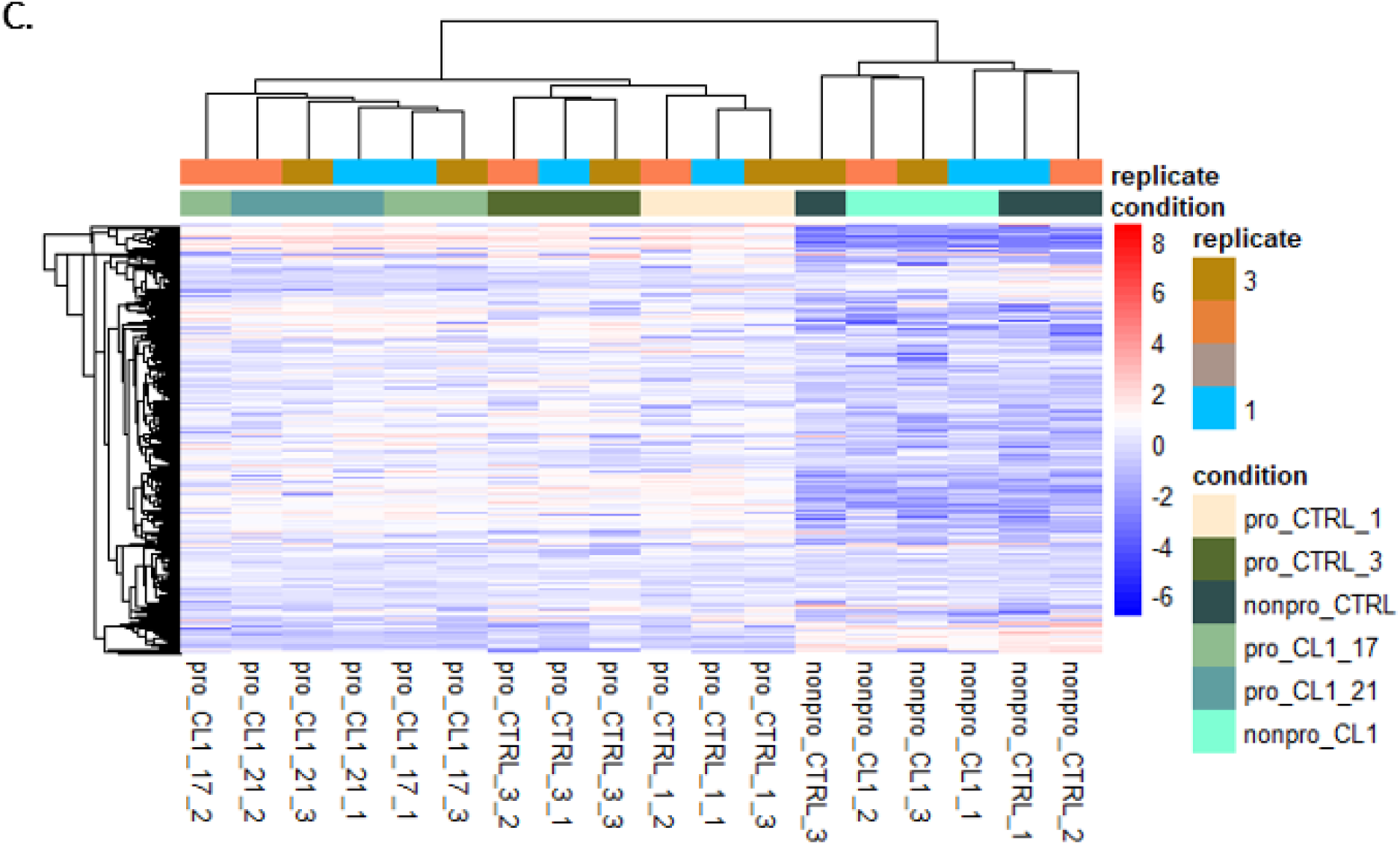
Pre-processing of the BAR proteomic analysis data in geCHO cell clones expressing E1E2. **(A)** Pre-processing of the BAR data involved plotting total proteins across all samples and their replicates. Nonproducer cell lines showed a lower number of proteins compared to the producer cell lines, as expected. Replicate samples contained relatively consistent protein numbers within their respective groups. **(B)** Principal Component Analysis (PCA) of the normalized proteomic dataset demonstrated clear separation between nonproducer (parental) cell lines and E1E2 producer clones. Principal component 1 (PCA1) captured 70% of the total variation, effectively distinguishing between the two cell line types. **(C)** Hierarchical clustering heatmap of the data revealed distinct separation between nonproducer and producer cell lines. Producer cell lines exhibited notably higher overall protein expression compared to nonproducer lines. The analysis identified specific protein groups with markedly different expression patterns: some proteins were highly expressed in producers but showed low expression in nonproducers, and conversely, some proteins were uniquely abundant in nonproducer cell lines.

### BAR identified PPIs of sE1E2 with host cell proteins

The processed BAR data were visualized with a heatmap showing that the nonproducers and the producers cluster separately (Figure 2C). Our analysis revealed that producer cell lines have proteins that are differentially expressed compared to the nonproducer cell lines. We observed distinct groups of proteins with higher expression in producer cells but low expression in nonproducer cells, and vice versa. Specifically, we identified 914 and 874 unique interactors in the CTRL and CL1 producer cell lines, respectively, that were detected only in these producer lines and not in their parental cell lines. We then performed Welch’s T-test comparing each producer cell line against its own parental cell line. This analysis identified 212 and 116 statistically significant interactors (Benjamini-Hochberg p-adjusted < 0.05 and Log2FC > 0) in CTRL and CL1 producer cell lines, respectively as illustrated in the volcano plots (Figure 3A). In total, we detected 974 and 903 unique potential protein interactors in CTRL and CL1 producer cell lines, respectively (Supplementary Table S1). Notably, 604 potential interactors were shared among all producer cell lines, suggesting a common set of proteins that may contribute to E1E2 production.

**Figure 3.**
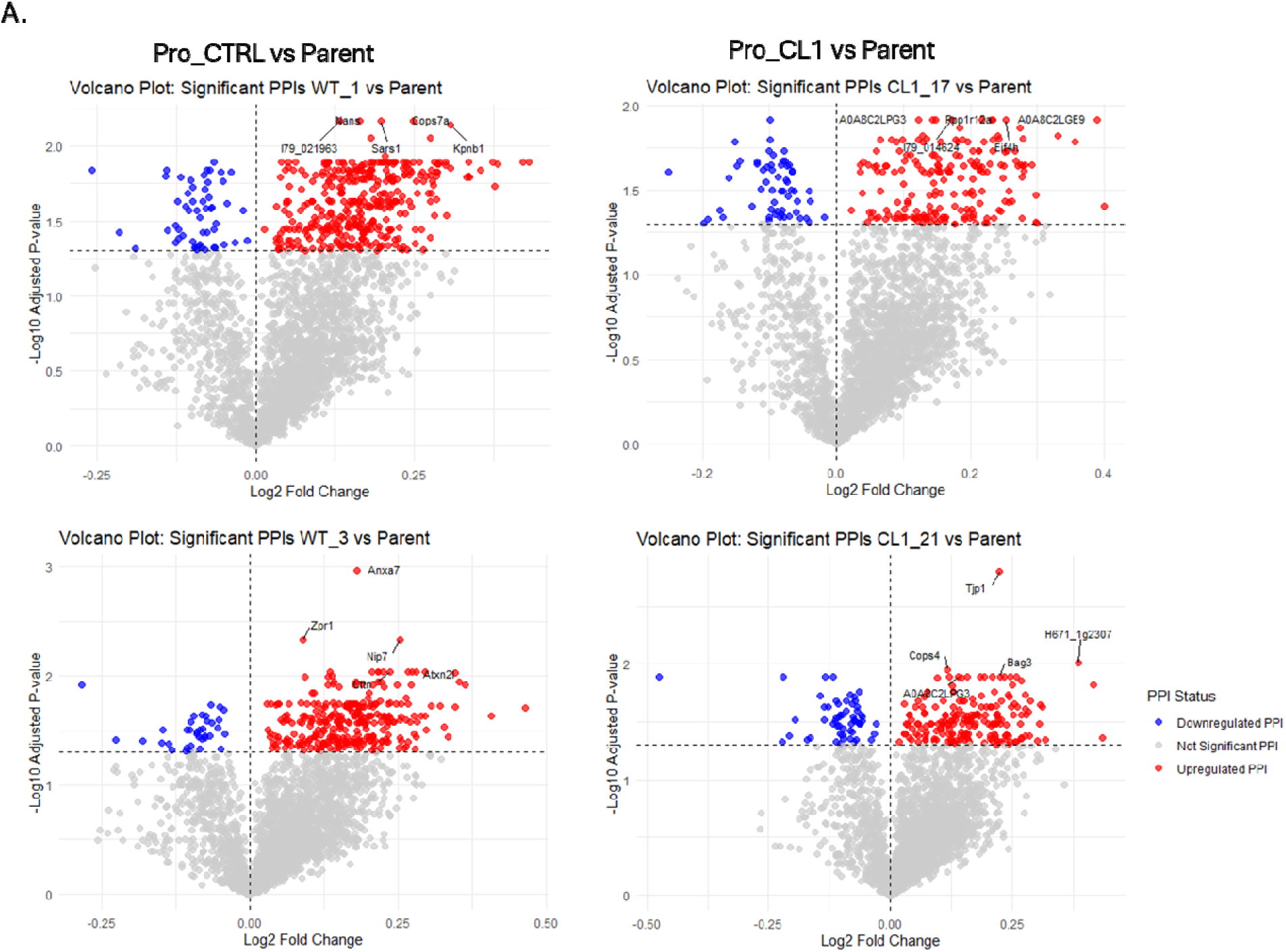
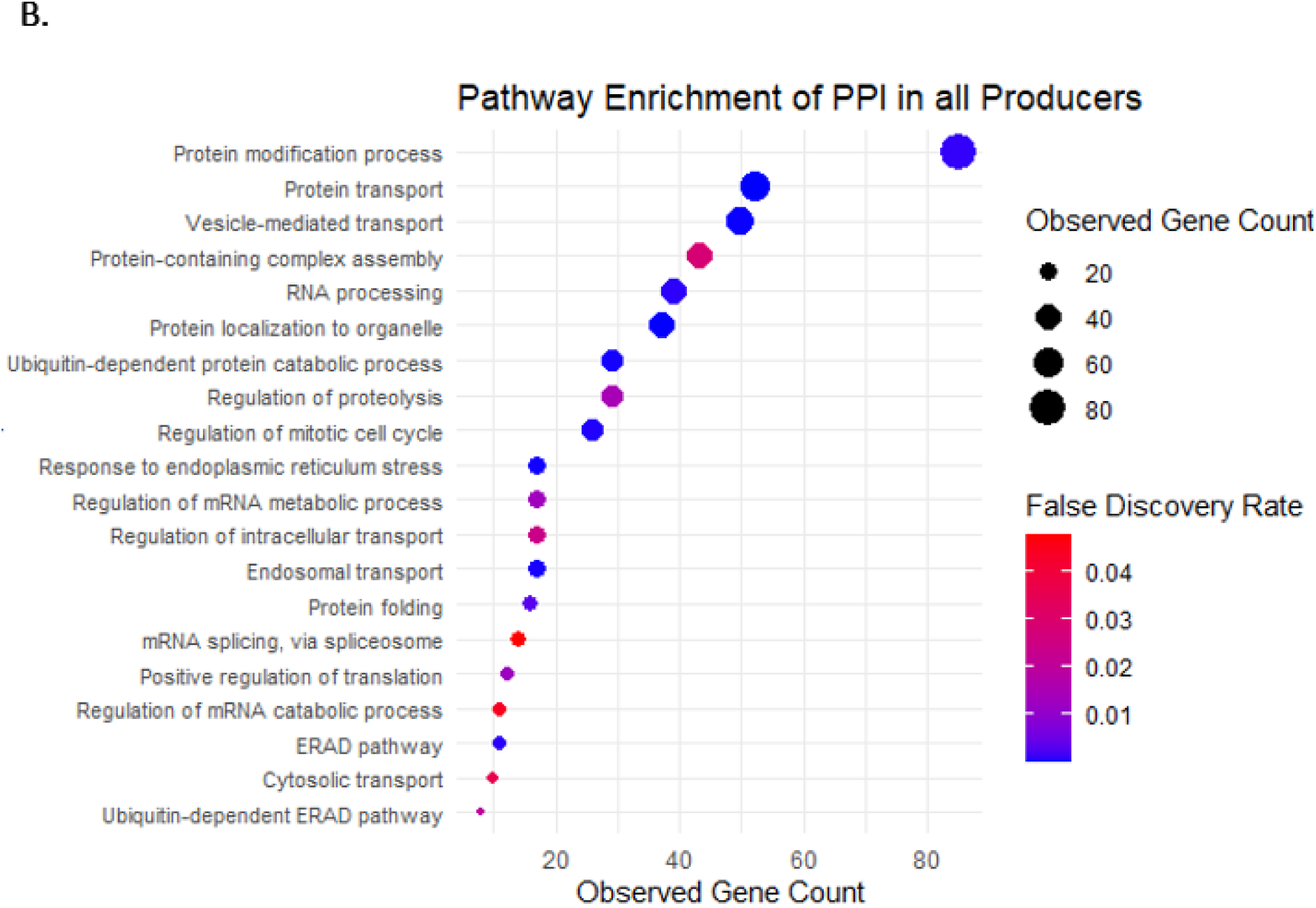
Identifying differentially expressed proteins in producer cells and pathway enrichment analysis. **(A)** Welch’s t-test was performed to compare proteomic data from BAR between producer and nonproducer cell lines, using respective parental cell lines as controls. Volcano plots visualize the statistical analysis, revealing proteins that are differentially expressed in the producer cell lines with statistical significance (padjusted < 0.05 and Log2FC > 0). **(B)** Gene enrichment analysis was performed on the 604 protein interactors that were either observed exclusively in the producer cell lines or differentially expressed, focusing on biological processes. The analysis revealed a comprehensive profile of cellular mechanisms, highlighting proteins critically involved in protein folding, intracellular transport, endoplasmic reticulum (ER) processes, and protein regulatory pathways.

Pathway enrichment analysis revealed that these PPIs are enriched in biological processes related to the secretory pathway, including protein transport, folding, and quality control. The analysis also captured proteins corresponding to RNA processing (mRNA splicing and translation) and mitotic cell cycle (Figure 3B). We identified 370 PPIs unique to CTL producer cell lines and 299 PPIs unique to CL1 producer cell lines. Both cell lines express PPIs related to cellular and metabolic processes, transport, and localization. Interestingly, the PPIs in CTL producer cell lines mainly correspond to RNA processing, mRNA/ncRNA metabolism, ribosome biogenesis, mitochondrion organization, and mitotic spindle processing. In contrast, PPIs in CL1 producer cell lines mainly correspond to regulation of cell death, apoptosis, programmed cell death, and small molecule metabolism.

### BAR and RNA-Seq analysis identifies potential protein interactors for improving sE1E2.SZ-H445P secretion

To refine our candidate gene list, we conducted differential gene expression analysis on bulk RNA-Seq data using DESeq2. We identified approximately 4,500 and 4,800 significant genes (p-adjusted < 0.05 and Log2FC > 0) in CTL and CL1 producers, respectively, (Figure 4A) when compared to their parental cell lines (Supplementary Table S2). Gene enrichment analysis using the enrichGO function in the clusterProfiler package in R was used to perform an over-representation analysis of the significantly upregulated genes in producing cells. It revealed enrichment in biological processes related to transport, DNA replication, and RNA processing (Figure 4B). Among the 604 shared PPIs, 151 genes were both significant (p-adjusted < 0.05) and upregulated, overlapping between proteomic and transcriptomic datasets. These proteins participate in various cellular processes, including cytoskeletal and motor functions, nuclear transport, transcription, ubiquitin pathways, transport mechanisms, ribosomal and translation-associated activities, signal transduction, cell communication, and mitotic cell cycle regulation.

**Figure 4.**
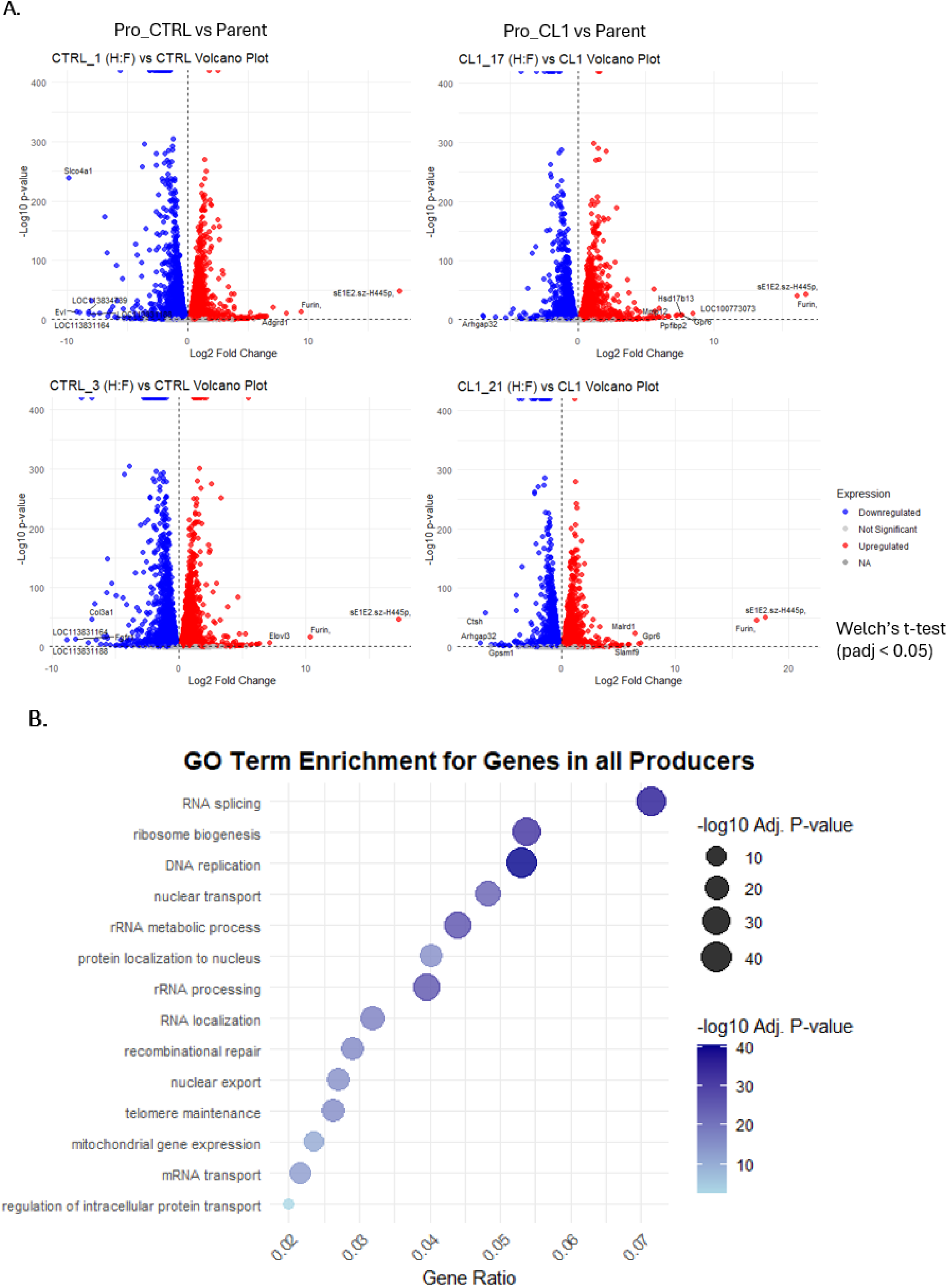
Transcriptomic analysis of bulk RNA-Seq in geCHO cells expressing sE1E2 **(A)** Bulk RNA-Seq was performed on geCHO cell clones expressing sE1E2, with respective parental cell lines serving as negative controls. Each cell line was analyzed in technical triplicates. Using the DESeq package in R and Welch’s t-test, we compared producer cell lines to their parental cell lines. The transcriptomic analysis revealed approximately 4,500 differentially expressed genes in CTRL and 4,800 differentially expressed genes in CL1 compared to their respective parental cell lines (padj < 0.05 and Log2FC > 0). **(B)** Gene Ontology enrichment analysis focused on biological processes across statistically significant genes in all producer cell lines. The analysis uncovered enriched pathways predominantly associated with cellular processes involving RNA metabolism, genetic material replication, and intracellular transport mechanisms.

We focused specifically on proteins involved in the secretory pathway, particularly related to proteostasis, ubiquitination, and transport mechanisms. Through our integrated proteomic and transcriptomic analysis, we narrowed our target list down to 30 genes associated with these processes. Based on statistical significance (p-adjusted < 0.05 and Log2FC > 0.2), we prioritized 12 high-confidence candidate genes for experimental validation.

### Overexpression of PPI proteins can increase sE1E2 secretion

We identified AP1M1, CUL4A, DNAJB1, IST1, MIA3, PSMA5, PSMD14, SPATA5, SUGT1, UFD1, YWHAH, and ZW10 as potential enhancers of the secretory pathway. These proteins participate in various cellular processes, including vesicular trafficking, protein degradation, and quality control, all of which are crucial for optimizing protein secretion efficiency.

To test their effects, we overexpressed each in CL1(H445P:Furin) #21, a CHO cell line expressing sE1E2, which has been associated with improved neutralization when producing sE2 (Kulakova et al., 2025). We transfected CL1(H445P:Furin) #21 cells in triplicate with the selected genes using Polyethylenimine (PEI) MAX transient transfection. RNA was extracted on day 2 post-transfection for complementary DNA (cDNA) synthesis and quantitative PCR (qPCR) analysis, which confirmed successful overexpression of all targeted genes (Figure 5A). On day 4 post-transfection, cell viability was assessed via cell counting, and supernatants were collected. The sE1E2 secretion was analyzed by western blotting, with band intensities quantified using ImageJ.

**Figure 5.**
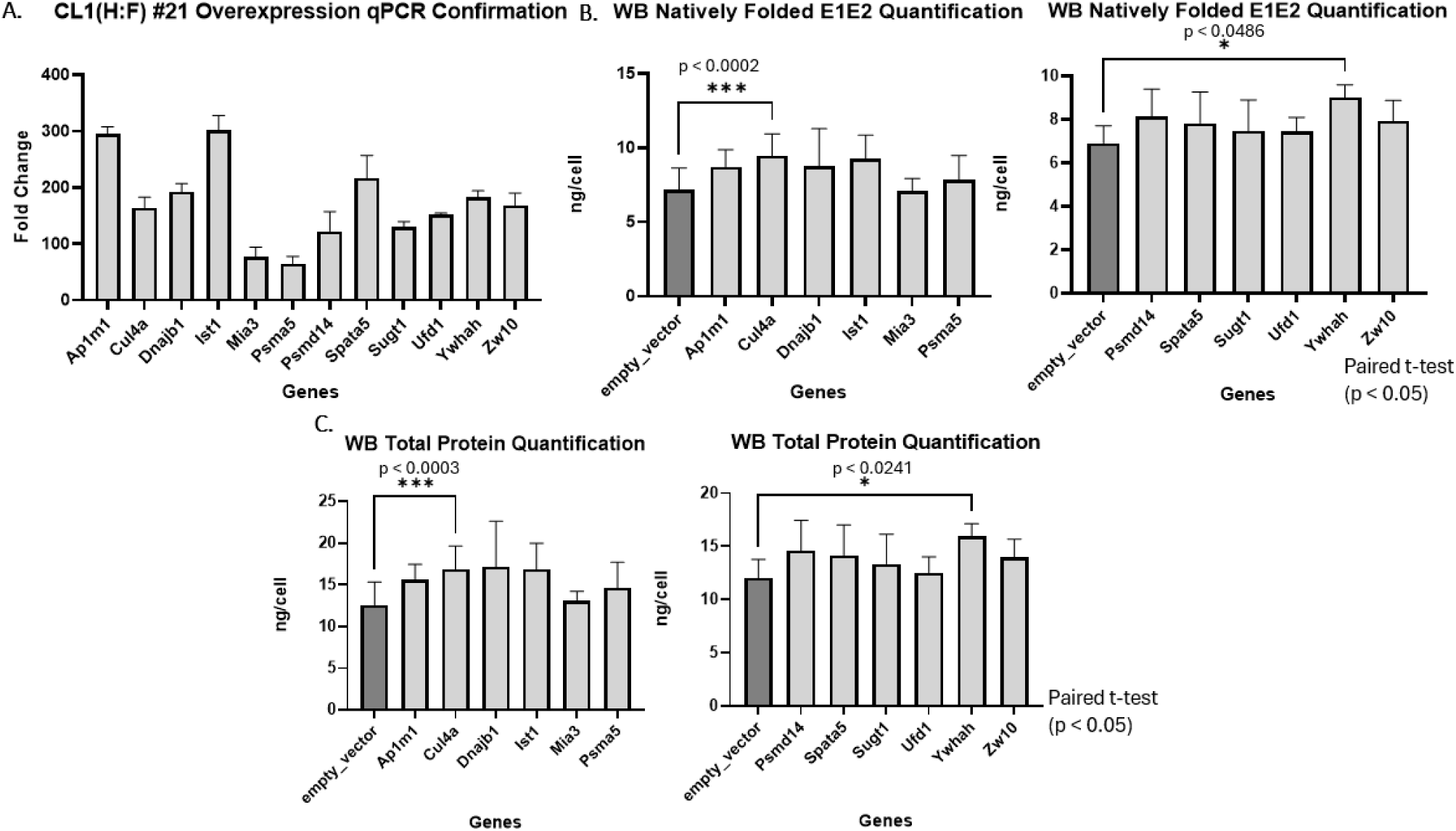
Validation of Candidate Genes Identified from Proteomic and Transcriptomic Datasets to Improve E1E2 Secretion. **(A)** Polyethylenimine (PEI) transient transfection was employed to overexpress genes selected based on proteomic and transcriptomic datasets. RNA was extracted on day 2 post-transfection and utilized for quantitative PCR (qPCR) to confirm gene overexpression. All selected candidate genes demonstrated successful overexpression following transient transfection. **(B)** Western blot analysis of the supernatant collected on day 4 post-transfection was performed to assess E1E2 secretion. Using ImageJ for quantification of natively folded sE1E2 bands, two genes, *Cul4a* and *Ywhah*, exhibited statistically significant (p < 0.05, paired t-test) improvements in the secretion of natively folded E1E2. **(C)** Total E1E2 protein secretion (encompassing both natively folded and uncleaved versions) was quantified using ImageJ. *Cul4a* and *Ywhah* demonstrated statistically significant (p < 0.05, paired t-test) enhancement in overall E1E2 protein secretion.

Based on viable cell counts and protein secretion analysis, CUL4A and YWHAH significantly enhanced the secretion of correctly folded sE1E2 (p < 0.05, paired t-test; Figure 5B) compared to the control (empty vector backbone) with fold changes of approximately 1.30 for both proteins. Additionally, overexpression of these genes significantly increased total sE1E2 protein secretion (p < 0.05, paired t-test), including both uncleaved and correctly folded forms, with similar fold changes of approximately 1.30 (Figure 5C). There was no significant difference in cell viability between the overexpressed genes and the control (p > 0.05, paired t-test; Supplementary Figure S1B). These two proteins were only detected in cells that expressed sE1E2 in the BAR proteomic data and not in the control cell lines, indicating their specificity to E1E2-expressing cells. The involvement of CUL4A and YWHAH in ubiquitination suggests that enhanced quality control checkpoints may benefit CHO cells in expressing HCV protein. Among all tested candidates, CUL4A and YWHAH emerged as the most promising for improving the secretion of both correctly folded and total sE1E2 protein.

## Discussion

The development of a sE1E2 protein vaccine by replacing the TMD with a synthetic scaffold, SZ, has shown promising results for scale-up and purification. While a secreted form of E1E2 could be produced in geCHO cells, the yield was too low for a viable vaccine. To enhance sE1E2 yield in geCHO cells, we analyzed the PPIs of sE1E2 using the BAR method to identify potential bottlenecks. We focused on the secretory pathway, which has previously been shown to enhance protein production through cellular engineering. For example, overexpression of calnexin and calreticulin, which are involved in the folding and quality control of glycoproteins, can increase thrombopoietin production by 1.9-fold (Chung et al., 2014). Similarly, overexpression of the transcription factor ATF4, which is involved in the unfolded protein response, can enhance IgG production (Haredy et al., 2013) and improve the production of recombinant human antithrombin III by twofold (Ohya et al., 2008). In our study, we identified two key proteins CUL4A and YWHAH, through the integrated analysis of BAR proteomic and RNA-Seq data; these two proteins significantly increased sE1E2 secretion when overexpressed.

CUL4A (Cullin 4A), a positive correlator with increased sE1E2 secretion, is a member of the Cullin-RING E3 ligases (CRLs) family. It forms the CRL4Cdt2 complex by interacting with Cdt2 (DNA replication licensing factor), a protein crucial for cell proliferation (Dar et al., 2014). This complex ubiquitinates proteins such as p21 (cyclin-dependent kinase inhibitor), Set8 (histone methyltransferase), Cdt1, and Cdc6 (cell division cycle 6) for degradation during S phase, ensuring cell cycle progression, DNA damage repair, prevention of re-replication, and proper replication initiation (Dar et al., 2014; Sharma & Nag, 2014). The 14-3-3ε and 14-3-3γ proteins positively regulate phosphorylated Cdt2 during the S phase, protecting it from degradation. In the absence of 14-3-3γ, destabilized Cdt2 can lead to increased levels of p21, Set8, and Cdt1, resulting in G2/M arrest (Dar et al., 2014).

Our second identified enhancer of sE1E2 secretion is YWHAH (14-3-3η), which belongs to the 14-3-3 protein family that binds to phosphoserine/phosphothreonine residues to regulate signal transduction and the cell cycle (Muslin et al., 1996). YWHAH, a tyrosine 3-monooxygenase/tryptophan 5-monooxygenase activation protein, promotes cell survival by inhibiting pro-apoptotic proteins such as Bad (BCL2 associated agonist of cell death), Bax (BCL2 associated X, apoptosis regulator), and ASK1 (Apoptosis signal-regulating kinase 1) (Nomura et al., 2003; Zha et al., 1996; Zhang et al., 1999). Previous studies have focused on engineering cells to improve cell viability and titer by knocking down pro-apoptosis genes such as BAX and BAK (Lim, et al. 2006). This provides a mechanistic explanation for our observed increase in protein production, as YWHAH naturally inhibits these same pro-apoptotic genes.

In addition, the 14-3-3 proteins have diverse cellular localizations and are found in the plasma membrane, ER, Golgi apparatus, nucleus, and mitochondria (Kongsamut & Eishingdrelo, 2023). Particularly, their presence in the ER and Golgi apparatus may directly influence the secretory pathway relevant to E1E2 processing. For example, 14-3-3 proteins interact with phosphorylated Nedd4-2, a HECT E3 ubiquitin ligase, inducing structural changes that may regulate substrate ubiquitination (Pohl et al., 2021).

Collectively, our findings suggest that CUL4A potentially positively regulates the cell cycle and proper replication initiation, while YWHAH likely positively influences cell viability by inhibiting proapoptotic proteins and interacting with the ubiquitin-proteasome system. Together, these proteins create a more favorable cellular environment for sE1E2 production and secretion. Additionally, simultaneously overexpressing both proteins would be of interest to determine whether they produce a significant increase in protein secretion compared to individual overexpression, since each protein targets a different pathway.

Despite identifying these two proteins that positively correlate with protein secretion, we still observed the secretion of the undesired uncleaved form of sE1E2 in the supernatant. Attempts to purify the correctly folded version for testing were unsuccessful due to the low yield of secreted E1E2. Future studies should investigate how sE1E2 is assembled intracellularly and the accessibility of the furin cleavage site in CHO cellular environments to improve cleavage efficiency. Enhancing cleavage efficiency would likely increase the proportion of correctly processed sE1E2, facilitating downstream purification and improving overall vaccine yield.

The BAR method provides a valuable approach for identifying CHO cell proteins that support sE1E2 secretion. However, since this method labels all proteins in proximity, it cannot distinguish between direct and indirect interactions and may include nonspecific interaction that can lead to false positives and negatives. To mitigate these limitations, we included (1) appropriate negative controls, (2) performed triplicate experiments to filter out noise, and (3) incorporated RNA-Seq as a mostly independent data type. This helped us to increase confidence in the identified PPIs.

Despite these challenges, the BAR method remains a valuable and user-friendly technique for studying both intracellular and secreted proteins, offering critical insights into the CHO secretory machinery. By identifying key components and mechanisms involved in protein secretion, it enhances our understanding of and provides a pathway to optimizing protein production. This approach demonstrates the potential of applying BAR to improve the secretion of other difficult-to-express proteins. When combined with other optimization strategies—such as plasmid design with enhanced promoters (Sou et al., 2023), site-specific integration using recombinase-mediated cassette exchange to prevent potential disruption of host genes (Shin et al., 2022), and bioprocess optimization (including fed-batch cultivation, pH control, aeration adjustment, media formulation, and temperature modulation) (Yang, 2019)—such cellular and process engineering approaches together create a comprehensive optimization regime that substantially enhances protein production.

## Materials and Methods

### Cell line engineering and protein expression

CHO-S cells (Life Technologies, Carlsbad, California, USA) were glycoengineered to create wt-like (CTRL) and CL1. The geCHO cells were generated using the gene editing technique CRISPR/Cas9, specifically using "CRISPy" to knock out *Sppl3* in CTRL cell line and *Sppl3*, *Mgat2*, *St3gal3/4/5*, and *B3gnt2* in CL1. The single cell cloning was done as described in Amann et al. paper (Amann et al., 2019).

### Cell culture

The geCHO cells were grown in 125 ml Erlenmeyer shaker flasks with baffled bottom (Fisher Scientific, USA) at 130 rpm in 37°C at 5% CO2 with maintenance media containing CD CHO Medium (Gibco, USA) supplemented with 8 mM L-Glutamine (Gibco) and 1x anti-clumping agent (Gibco). The geCHO cells expressing E1E2 and furin were grown in media containing 1 mg/ml Geneticin Selective Antibiotic (G418 sulfate; Gibco) and 0.75 mg/ml Hygromycin B (Gibco) for selection.

### CellTiter-Blue Assay-Kill Curve

CellTiter-Blue assay was used to measure the viability of CHO-S cell lines to determine the concentration of Geneticin Selective Antibiotic (Gibco) and Hygromycin B (Gibco) to be used for selection after transfection. The cells were grown in 125 ml shake flasks in maintenance media at 5% CO2 and were > 95% viable before the experiment was conducted. Cells were seeded at 0.2 × 10^6^ cells/mL in non-treated 12-well plates (Greiner Bio-One, Kremsmunster, Austria) and incubated with different concentrations (0-1 mg/mL final concentration) of G418 or Hygromycin B at a final volume of 1 ml. Each concentration was measured in technical duplicates. The assay was conducted for 10 days and the media was changed every 2 days. To measure viability and cell count, 100 μL was taken every other day from each well and mixed with 20 μL of the CellTiter-Blue reagent in a 96 well plate (Sarstedt, Numbrecht, Germany). The reaction was incubated at 37°C for 2 hours. Then the plate was read using the BioTek Synergy MX microplate reader (Agilent, Santa Clara, CA, USA) with excitation wavelength at 530 nm and emission wavelength at 590 nm. The plate was shaken at medium speed for 1 minute before reading.

### Plasmid Design

The sE1E2.SZ plasmid was generated as described previously (Metcalf et al., 2023) with a mutation in Domain D in E2 forming sE1E2.SZ-H445P (Pierce et al., 2020), and the furin plasmid was generated as described previously (Guest et al., 2021). The furin plasmid was redesigned with a new backbone, pcDNA4-TO-Hygromycin-sfGFP-MAP purchased from Addgene. It contains a CMV promoter expressing the gene for furin of 2385 bp with a bGH poly(A) signal, an ampicillin resistance marker and a hygromycin resistance marker for post-transfection selection.

### Stable Transfection

The geCHO cells were initially transfected with the furin plasmid and cloned. The cloned furin cell lines were then transfected with sE1E2.SZ-H445P plasmid and cloned again. At 24 hours before transfection, the cells were passaged at 0.8-1.0 × 10^6^ cells/mL with CD CHO Medium (Gibco) supplemented with 8 mM L-Glutamine (Gibco). On the day of transfection, the cells were filtered with a 40 μm nylon mesh cell strainer (Falcon) before cell count. Twenty-five million cells were harvested, which was enough to transfect one negative control and two samples with plasmid. Cells were spun down at 200g for 4 minutes and washed once with warm PBS. Cells were resuspended in Buffer R from the Neon Transfection System 100 μL Kit (Invitrogen) with 12 μg of furin plasmid. Subsequently, DNA/cell suspension containing 5 × 10^6^ cells for each transfection were transfected using the Neon Transfection System with the following parameters: Voltage: 1550 V, pulse width: 10 ms, pulse number: 2 pulses. The transfected cells were transferred to a 6-well plate (GenClone; Genesee Scientific, El Cajon, CA, USA) with pre-warmed mixture media (2 mL/well). The plate(s) were placed on a shaker at 90 rpm at 37°C with 5% CO2. The next day, the viability of the cells was checked and the shaker speed was increased to 120 rpm. On day 2, the cells were expanded to a shake flask in media containing Hygromycin B. When the cells expanded, cells were passaged at 0.2 × 10^6^ cells/mL. The total cell count and viability were measured every other day, and media was changed every 3 days until the cells recovered and had > 95% viability. Afterward, cells were harvested for western blot for confirmation. The stable polyclonal cells were then cloned as described and the stable furin clones were transfected with 12 μg of sE1E2.SZ-H445P plasmid to generate stable clones expressing sE1E2.SZ-H445P and furin.

### Cloning stable polyclonal cell lines

To form a stable cell line, cloning was done for each transfection. Cloning media contained CD CHO Medium (Gibco), 80% Ex-Cell CHO Cloning Medium (Sigma-Aldrich, St Louis, MO, USA), 8 mM L-Glutamine (Gibco), 1× ClonalCell-CHO ACF Supplement (Stemcell Technologies, Vancouver, Canada), 1 mg/mL Geneticin Selective Antibiotic (G418 sulfate; Gibco) and 0.75 mg/mL Hygromycin B (Gibco). Cells needed to be > 95% viable to be used for cloning. Cells were serially diluted at 1:10 dilution in media with 1 mg/mL of G418 and 0.75 mg/mL Hygromycin B containing no anti-clumping until the cells were at 1,000 cells/mL in 15 mL falcon tubes at a volume of 10 mL. Non-treated 96-well plates for suspension (N=3) (Genesee Scientific) were filled with 180 μL of cloning media into each well except column 1. Column 1 was plated with 200 μL of the 1,000 cells/mL suspension. Then the cells were serially diluted down to 10 cells (column 3) by taking 20 μL from previous wells. Onward, 20 μL was taken from the 10 cells well (column 3). This gave 1 cell/well from column 4-12. The plates were incubated at 37°C in 5% CO2 with no shaking. After 2 weeks, the plates were checked for single clones under microscopy. After 3 weeks, cells were expanded to non-coated 24-well plates (Genesee Scientific) prefilled with 400 μL of maintenance media containing antibiotics with no shaking. Cells were grown for a week before transferring to a non-coated 12-well plate (Genesee Scientific) prefilled with 600 μL of maintenance media containing antibiotics with shaking. After ∼1 week or when cells expanded, half of the cells were taken for dot blot while the other half was expanded to 3 mL in a 6-well non-coated plate (Genesee Scientific) shaken at 130 rpm. Viability was performed on the cells once they grew, and they were expanded to 125 mL shake flasks.

### Dot blot for screening clones

To screen the clones in 12-well plates, dot blot was used to ensure furin expression in lysate and secretion of E1E2 in the supernatant of these clones before further expansion. Cells and supernatants were taken in 12-well plates. The cells were lysed using 1× RIPA buffer (Thermo Fisher Scientific, USA), 1× cOmplete Protease Inhibitor Cocktail (Roche, Basel, Switzerland) in distilled water. The 0.2 μm Nitrocellulose membrane (BioRad) was loaded with 10 μL of lysate or supernatant. The membrane(s) were dried for ∼1 hour, and then blocked for 1 hour in the Intercept Blocking Buffer (LI-COR) in a rocker at room temperature. Primary antibodies used were: 1:2,000 Furin Monoclonal antibody mouse (67481-1-Ig; Proteintech, Rosemont, IL, USA); 1:500 E2 (HCV1); or 1:400 HepC E1 Mouse Monoclonal IgG 100 μg/mL (LO518, Santa Cruz Biotechnology, Dallas, TX, CA). The primary antibody was diluted in Intercept antibody diluent (LI-COR, Lincoln, NE, USA) and stained overnight in the rocker at 4°C. The membrane was washed three times with 0.1% TBST for 10 minutes each. Secondary antibody was diluted in the Intercept Dilution Buffer (LI-COR): 1:15,000 IRDye 800CW Goat anti-Human IgG Secondary Antibody (LI-COR) or 1:15,000 IRDye 680RD Goat anti-Mouse IgG Secondary Antibody to the membrane. It was incubated for 1 hour at room temperature in the rocker. Then it was washed with 0.1% TBST 3× for 10 minutes each. Then the membrane was imaged using the LI-COR 9120 Odyssey Infrared Imaging System (LI-COR).

### Western-blot for furin and E1E2 transfection

Western blot was performed to analyze protein secretion of E1E2 using a precast polyacrylamide gel, 4-15% Mini-Protean TGX Gels (Bio-Rad, Hercules, CA, USA). The samples were run under reducing conditions by mixing with loading dye (4x Laemli + 10% β-Mercaptoethanol) (Bio-Rad) and incubating at 95°C for 5 minutes. The gel was run on a Mini-PROTEAN Tetra Vertical Electrophoresis Cell (Bio-Rad) at 160V for 40 minutes with the Precision Plus Protein standards ladder (Bio-Rad). The gel was transblotted onto 0.2 μm nitrocellulose membranes (Bio-Rad) with the Trans-Blot Turbo Transfer System (Bio-Rad). Primary antibodies used were: 1:2,000 Furin mouse Monoclonal antibody (67481-1-Ig; Proteintech); 1:1000 Anti-HCV E2 mouse, clone AP33 25 MG (MABF2820; Sigma-Aldrich); or 1:400 HepC E1 Mouse Monoclonal IgG 100 μg/mL (LO518; Santa Cruz Biotechnology). The primary antibody was diluted in Intercept antibody diluent (LI-COR) and stained overnight in the rocker at 4°C. The membranes were blocked with Intercept Blocking Buffer (LI-COR) for 1 hour. Rituximab was detected with IRDye 800CW Goat anti-human IgG Secondary Antibody (LI-COR) at 1:15,000 in Intercept Antibody Diluent (LI-COR) for 1.5 hours at room temperature. The membranes were washed 3x at 10 minutes with 0.1% TBST and imaged using the LI-COR 9120 Odyssey Infrared Imaging System (LI-COR).

For staining of intracellular biotinylated proteins, 20 μg of total protein from labeled cells was loaded and transblotted. The membrane was blocked by 3% BSA in 0.1% TBST for 1 hour and probed with HRP-conjugated streptavidin (Abcam, Cambridge, UK) diluted in blocking buffer at 1:2000 for 40 minutes. For visualizing the proteins’ bands, the Clarity Western ECL Substrate (Bio-Rad) was used.

### Quantify PPIs of E1E2 in geCHO cells using the BAR method

BAR was utilized to target sE1E2 in 2 clones from each geCHO cell lines expressing sE1E2 and identify its PPIs in each clone. To do this, the geCHO clones expressing E1E2 with the non-producing geCHO cells as control, each done in technical triplicates were harvested at the mid-exponential phase, fixed with 4% PFA in PBS (Thermo Fisher Scientific), and permeabilized with 0.4% PBST. Peroxidases were inactivated with 0.4% H_2_O_2_ (Fisher Scientific) and cells were blocked with 5% goat serum (Gibco) in 1% BSA in PBST. The cells were then incubated with E2 (HCV1, human) in blocking buffers overnight with rotation at 4°C. Next day, the samples were incubated with Goat anti-human IgG HRP (Invitrogen) for 1 hour at room temperature with rotation. Proximal protein biotinylation occurred with treatment of H_2_O_2_ and tyramide-biotin using the TSA Biotin Reagent Pack (SAT700001EA; Akoya Biosciences, Marlborough, MA, USA), resulting in tyramide-biotin radicalization and deposition onto proximal proteins. The reaction was stopped with 0.5 M sodium ascorbate in PBS (Sigma-Aldrich). Lysate was extracted using 1.5% SDS (G Biosciences, St Louis, MO) and 1% Sodium Deoxycholate (bioWorld, Dublin, OH, USA) in 0.1% PBST and heated at 99°C for 1 hour with mild shaking. The samples were spun down at max speed for 5 minutes, and supernatant was taken for quantification using BCA Protein Assay (Lamda Biotech, Inc., St Louis, MO, USA). The samples were stored at -80°C until being sent to Sanford Burnham Prebys Proteomic Core (San Diego, CA, USA) for LC-MS/MS.

### LC/MS-MS on biotinylated samples

The BAR samples were sent to Sanford Burnham Prebys Proteomic Core for LC-MS/MS analysis. Biotinylated proteins were affinity-purified using the Bravo AssayMap platform (Agilent) with AssayMap streptavidin cartridges (Agilent). Briefly, cartridges were first primed with 50 mM ammonium bicarbonate, and then proteins were slowly loaded onto the streptavidin cartridge. Background contamination was removed with 8M urea, 50 mM ammonium bicarbonate. Finally, cartridges were washed with Rapid digestion buffer (Promega, Rapid digestion buffer kit) and on-cartridge digestion of the bound proteins was performed using mass spectrometry-grade Trypsin/Lys-C Rapid digestion enzyme (Promega, Madison, WI) at 70°C for 1 hour. The resulting peptides were desalted on the Bravo platform using AssayMap C18 cartridges. Organic solvents were removed using a SpeedVac concentrator prior to LC-MS/MS analysis. The dried peptides were reconstituted in 2% acetonitrile with 0.1% formic acid and quantified using a NanoDropTM spectrophotometer at A280 nm (Thermo Fisher Scientific). Samples were analyzed via LC-MS/MS using a Proxeon EASY nanoLC system (Thermo Fisher Scientific) coupled with a Q-Exactive Plus mass spectrometer (Thermo Fisher Scientific).

Peptide separation was conducted on an analytical C18 Aurora column (75µm x 250 mm, 1.6µm particles; IonOpticks) at a flow rate of 300 nL/min. The 75-minute gradient applied was: 2% to 6% B in 1 minute, 6% to 23% B in 45 minutes, 23% to 34% B in 28 minutes, and 34% to 48% B in 1 minute (A: 0.1% FA; B: 80% ACN with 0.1% FA). The mass spectrometer was operated in positive data-dependent acquisition mode. MS1 spectra were measured in the Orbitrap in a mass-to-charge (m/z) of 375 – 1500 with a resolution of 60,000. Automatic gain control target was set to 4 x 10^5 with a maximum injection time of 50 ms. The instrument was set to run in top speed mode with 1-second cycles for the survey and the MS/MS scans. After a survey scan, the most abundant precursors (with charge state between +2 and +7) were isolated in the quadrupole with an isolation window of 0.7 m/z and fragmented with HCD at 30% normalized collision energy. Fragmented precursors were detected in the ion trap as rapid scan mode with automatic gain control target set to 1 x 10^4^ and a maximum injection time set at 35 ms. The dynamic exclusion was set to 20 seconds with a 10 ppm mass tolerance around the precursor.

Mass spectra were analyzed using MaxQuant software (Tyanova, Temu, & Cox, 2016), version 1.5.5.1. MS/MS spectra were searched against the Cricetulus griseus Uniprot and trEMBL protein sequence database (downloaded in September 2024) and GPM cRAP sequences (common protein contaminants).

Precursor mass tolerance was set to 20 ppm for the initial search, which included mass recalibration, and 4.5 ppm for the main search. Product ions were searched with a mass tolerance of 0.5 Da. The maximum precursor ion charge state for the search was set to 7. The enzyme specificity was set to trypsin, allowing up to two missed cleavages.The target-decoy-based false discovery rate (FDR) filter for spectrum and protein identification was set to 1%.

### BAR analysis

The BAR proteomic mass spectrometry dataset was preprocessed and analyzed using the DEP package in R Studio to identify differentially expressed proteins. Proteins were assigned unique names, and samples were filtered to remove proteins with excessive missing values. Specifically, proteins were retained only if they were identified in at least 2 out of 3 replicates for any given condition. Samples were normalized using variance stabilizing transformation (VSN), and missing values were imputed based on data missing not at random (MNAR) using a manually defined left-shifted Gaussian distribution. The data were then visualized using Principal Component Analysis (PCA) of the normalized and imputed data. Proteins were filtered using the following criteria: detected in ⅔ replicates for each producer clone, present in < 2 replicates in control, or present in ⅔ replicates in control but having a fold change > 0 and adjusted p-value (p-adjusted) < 0.05. These were marked as potential hit proteins. Welch’s T-test was performed comparing producer cell lines versus non-producer cell lines to identify proteins that were significantly expressed with p-adjusted < 0.05 and fold change of > 0 in the producer cell lines.

### RNA-Seq prep and analysis

Each of the geCHO cell clones expressing sE1E2 along with its parental cell line, each in triplicates, were harvested during the same time as the sample used for BAR. Total RNA was extracted from each sample using the RNeasy Kit (Qiagen, Hilden, Germany) following the manufacturer’s instructions and quantified using nanodrop at A260 nm. Samples were sent to UCSD IGM Genomics Center for mRNA library preparation and sequencing. RNA quality was analyzed using the Agilent Tapestation 4200, and only samples with an RNA Integrity Number (RIN) exceeding 8.0 were selected for library construction. Libraries were prepared using the Illumina Stranded mRNA Sample Prep Kit with Illumina RNA UD Indexes (Illumina, San Diego, CA), following the manufacturer’s guidelines. The finalized libraries were multiplexed and sequenced on an Illumina NovaSeq X Plus platform, generating 150 base pair (bp) paired-end reads (PE150) with an approximate depth of 25 million reads per sample. The samples were demultiplexd with bcl2fastq v2.20 Conversion Software. Adapter sequence trimming and quality control (FastQC) of the sequences were performed using trimgalore. Reads were mapped to the CHO GCF_003668045.3 CriGri-PICRH-1.0 genome, aligned and quantified with kallisto. The samples are analyzed using R studio using the DEseq2 package.

### Generating plasmids for overexpression

The Ap1m1, Dnajb1, Cul4a, Ist1, Mia3, Psmd5, Psmd14, Spata5, Sugt1, Udf1, Ywhah, Zw10, and the empty vector (pcDNA3.1(+)) plasmids were designed from Chinese hamster (Cricetulus griseus) and purchased from GenScript (Piscataway, NJ, USA). The plasmids were transformed in DH5alpha Competent Cells (Invitrogen, USA) and expanded in Miller’s LB Broth (Corning Inc., Corning, NY, USA) supplemented with 100 μg/ml of ampicillin (Sigma-Aldrich). The plasmids were extracted using QIAprep Spin Miniprep Kit (Qiagen).

### Transient Transfection

Polyethylenimine (PEI) Max transient transfection was used for overexpression of the validation genes. PEI MAX-Transfection Grade Linear Polyethylene Hydrochloride (MW 40,000) (Kyfora Bio, Warrington, PA, USA) was dissolved at 1 mg/ml in distilled water and pH was adjusted to 7.0 using 1M of NaOH. The dissolved PEI was sterile filtered in 0.22 μm steritop (MilliporeSigma, Burlington, MA, USA). A day before transfection, cells were passaged at 0.5-0.8 × 10^6^ cells/ml in media containing no anti-clumping. On the day of transfection, cell counts were done and were at least > 95% viable for transfection. Cells were harvested and resuspended at 1.0 × 10^6^ cells/ml in media containing no anti-clumping. Cells were transfected with 1 μg plasmid and 7 μg of PEI Max solution per ml of cell culture (5% of total culture volume). The DNA/PEI mixture was incubated in OptiPRO SFM (Life Technologies) for 20 minutes at room temperature before adding to the cells in 6-well non-treated, polystyrene, flat bottom wells plate (Genesee Scientific). The cells were shaken at 130 RPM at 37°C in 5% of CO2. The day post transfection, 1× of anticlumping agent (Gibco) and 0.6 mM valproic acid sodium salt (Sigma-Aldrich) was added to each well. Some cells were harvested at day 2 post transfection for RNA extraction. Viability was performed on day 4 post transfection and supernatant was harvested.

### qPCR to confirm gene overexpression

RNA extraction was done following the RNeasy Mini kit protocol (Qiagen). RNA was converted to cDNA following the SuperScript II RT protocol (Thermo Fisher Scientific) with 1 μg of RNA and RNAseOUT Recombinant Ribonuclease Inhibitor (Life Technologies). qPCR was performed using iTaq™ Universal SYBR Green Supermix (Bio-Rad) using primers targeting the validating genes purchased from IDT DNA (USA) (Table 1) with Gnb1 as the housekeeping gene in 96-well PCR Plates (Bio-Rad). The samples were run as follows: 95°C for 2 min; 40×: 95°C for 10 sec, 60°C for 30 sec; 65°C for 5 sec, 95°C for 5 sec. Using the ΔΔCt method, the relative expression levels of each gene were calculated by normalization to the expression level of the normalization gene. Each experiment included no-template controls and was performed using technical duplicates.

**Table 1:**
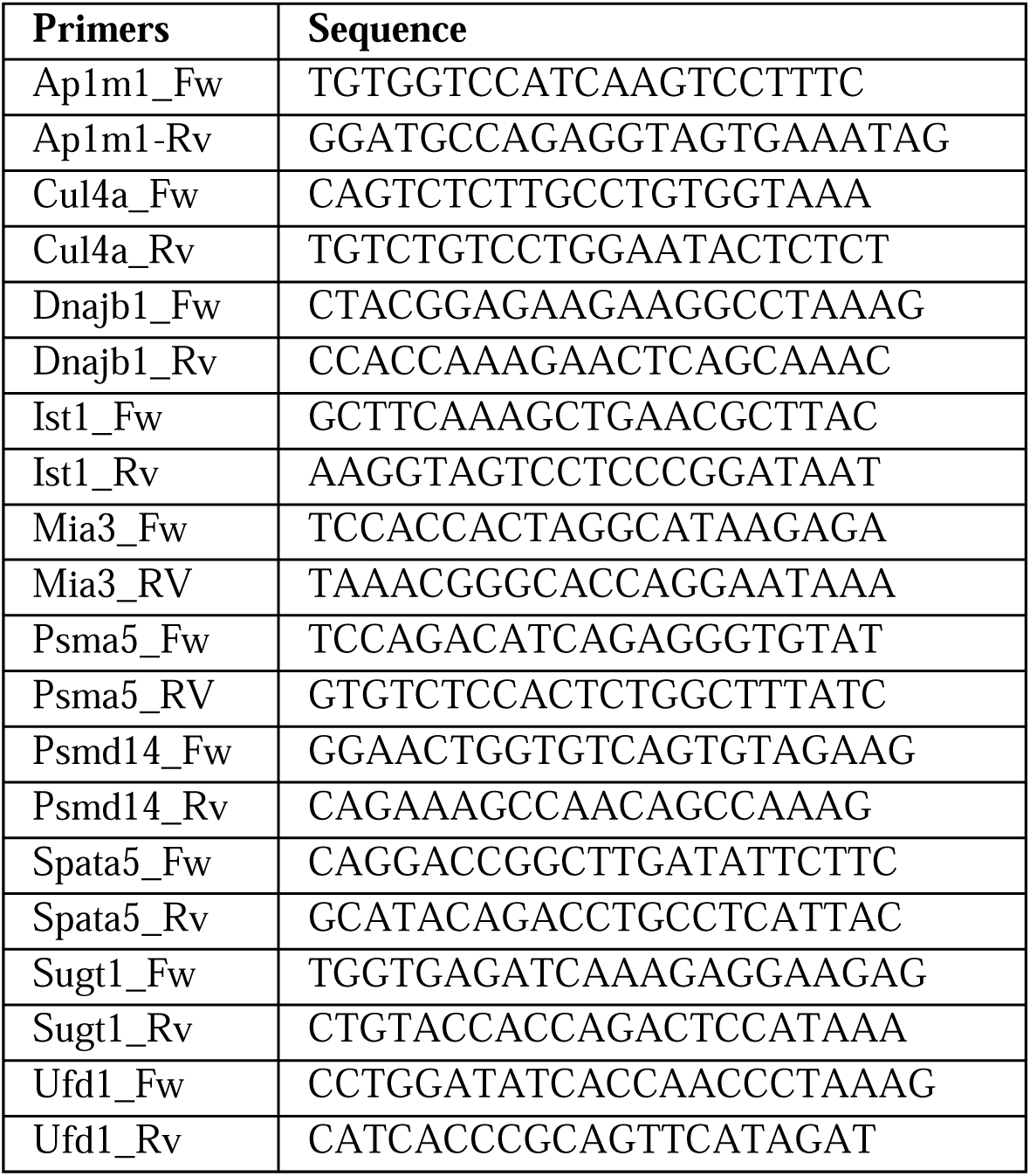
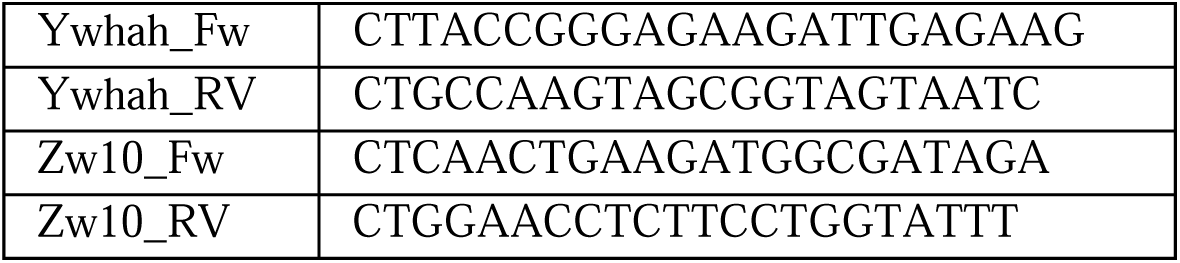
Primers of validating genes for qPCR.

### Statistical Analyses

Statistical analysis of BAR proteomic data was performed using the DEP package (version 1.24.0) in RStudio (version 4.3.3) and Microsoft Excel. RNA sequencing data was analyzed using the DESeq2 package (version 1.42.1) and clusterProfiler package (version 4.10.1) in RStudio, with genome annotation provided by the org.Hs.eg.db package (version 3.18.0). Data visualization included volcano plots, RNA-Seq PCA plots, and enrichment pathway plots, which were generated using the ggplot2 package (version 3.5.0). BAR proteomic PCA plots were created with the DEP package. Bar graphs for qPCR and validated gene expression data were generated using GraphPad Prism using Paired T-test (version 10.4.0). All figures were designed and assembled using BioRender.com.

## Acknowledgments

This work was supported by generous funding from NIH (R35 GM119850, R01 AI132212, R01 AI168048), NSF (CBET-2030039), and the Novo Nordisk Foundation (NNF20SA0066621).

## Declaration of Interests

NEL is a scientific advisor for CHO Plus, and cofounder of NeuImmune, Inc. along with Thomas R. Fuerst, and Augment Biologics, Inc.

## Data availability

Study findings are supported by data available upon request from the corresponding author. Mass spectrometry proteomics data, raw and processed, has been uploaded to the MassIVE (member of the proteome Xchange consortium) repository at https://doi.org/10.25345/C5ZZ0V, reference number PXD063086. Raw and processed transcriptomic data has been uploaded to GEO with the data set identifier [GSE294982].

