## Supplementary Figure S1 for "Enhanced Production of HCV E1E2 Subunit Vaccine Candidates via Protein-Protein Interaction Identification in Glycoengineered CHO cells"

**Supplementary Figures:**

**A.**

CTRL (H:F) #1

**250 -**

**150 -**

**100 -**

**75 -**

**50 -**

**37 -**

**25 -**

**20 -**

**15 -**

**10 -**

**kDa**

300 ng sz.E1E2 lysate

Day1 Sup

Day2 Sup

Day4 Sup

Day3 Sup

Day4 lysate

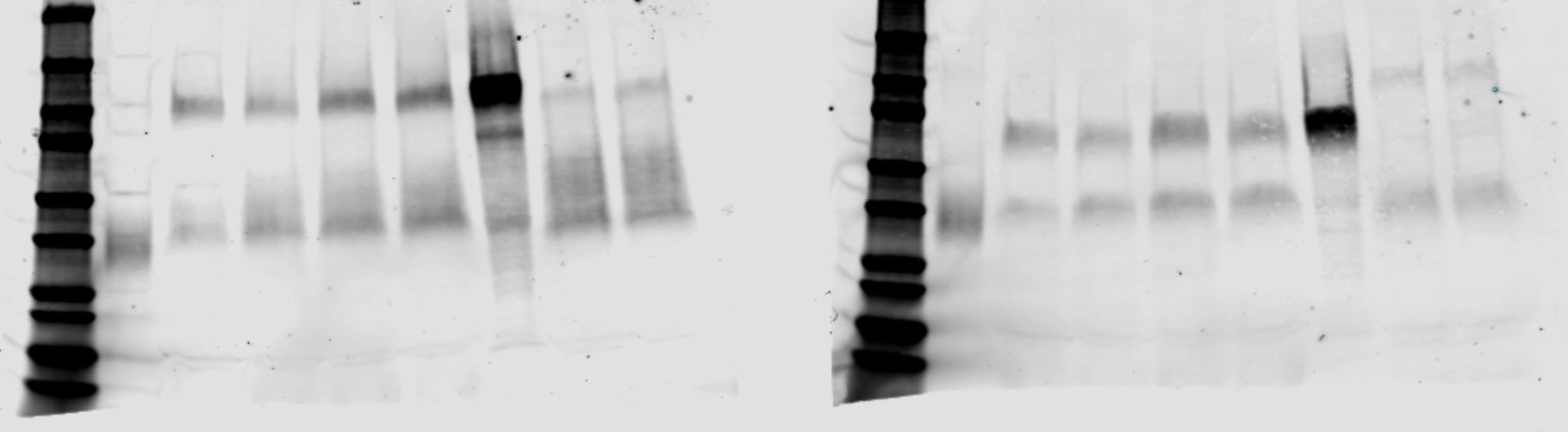

CTRL (H:F) #3

**250 -**

**150 -**

**100 -**

**75 -**

**50 -**

**37 -**

**25 -**

**20 -**

**15 -**

**10 -**

**kDa**

300 ng sz.E1E2 lysate

Day1 Sup

Day2 Sup

Day4 Sup

Day3 Sup

Day4 lysate

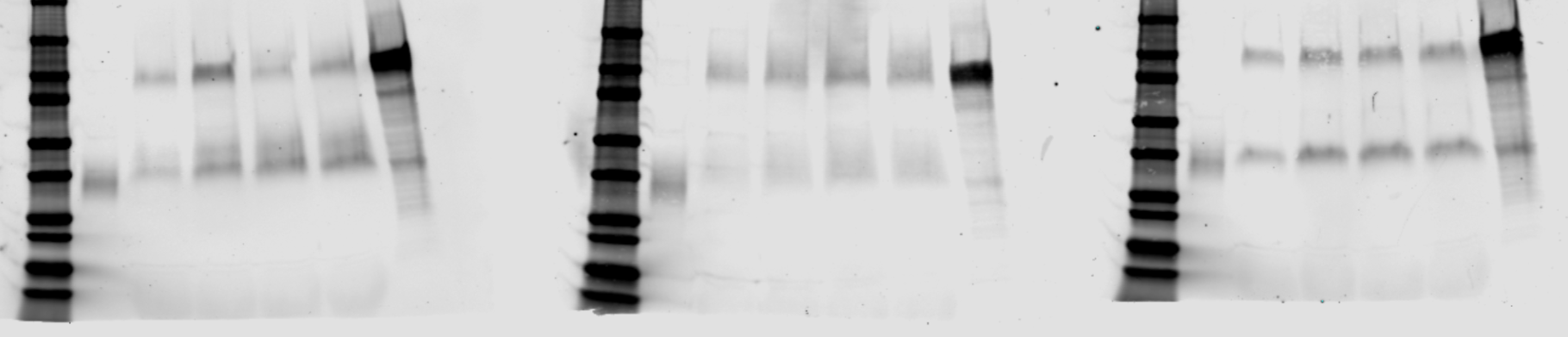

CL1(H:F) #17

**250 -**

**150 -**

**100 -**

**75 -**

**50 -**

**37 -**

**25 -**

**20 -**

**15 -**

**10 -**

**kDa**

300 ng sz.E1E2 lysate

Day1 Sup

Day2 Sup

Day4 Sup

Day3 Sup

Day7 lysate

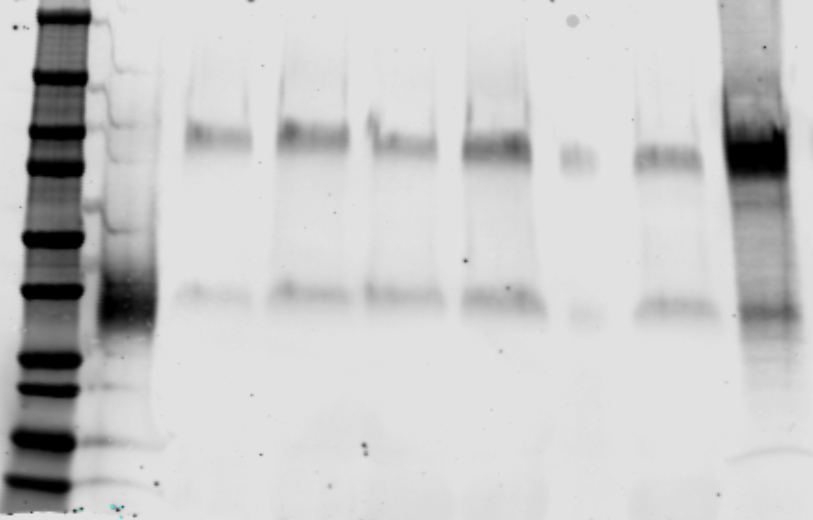

Day7 Sup

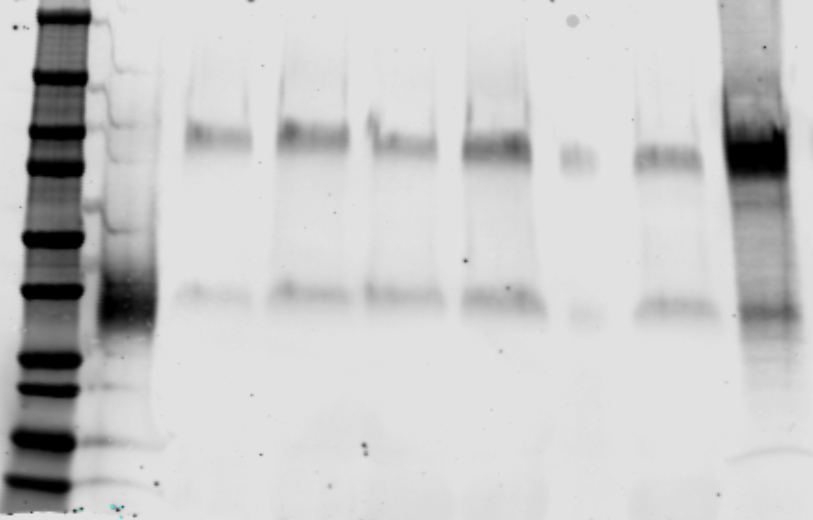

**< uncut E1E2 (94 kDa)**

**< E1 (~30 kDa)**

**250 -**

**150 -**

**100 -**

**75 -**

**50 -**

**37 -**

**25 -**

**20 -**

**15 -**

**10 -**

**kDa**

300 ng sz.E1E2 lysate

Day1 Sup

Day2

Sup

Day4

Sup

Day3 Sup

Day4

lysate

CL1(H:F) #21

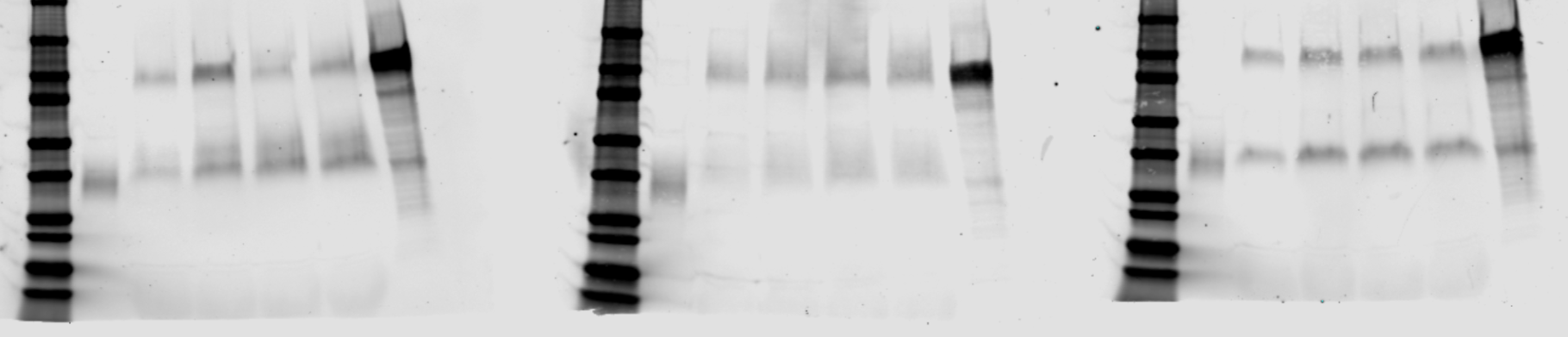

**< uncut E1E2 (94 kDa)**

**< E1 (~30 kDa)**

**B.**

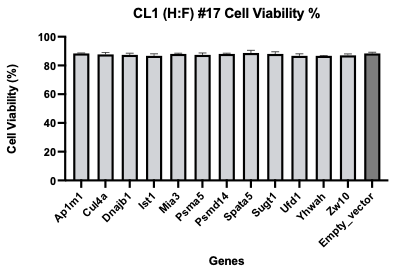

**Supplementary Figure S1. Expressing and confirming sE1E2.SZ-H445P in geCHO cell clones.**
**(A)** Stable clones expressing E1E2 were established, and two clones from each geCHO cell line were selected for biotinylation by antibody recognition (BAR) analysis. The selected clones comprised CTRL (H:F) #1, CRTL (H:F) #3, CL1 (H:F) #17, and CL1 (H:F) #21. Reduced western blot analysis was performed for each clone, with supernatant collected over a 4-day period and cell lysate collected on day 4. The nitrocellulose membrane was probed with an anti-E1 primary antibody (mouse, Santa Cruz Biotechnology) at a 1:400 dilution, followed by a secondary antibody (goat anti-mouse IR 680, LI-COR Biosciences). Membrane imaging was conducted using a LI-COR imaging system. The expected bands for furin-cleaved E1E2 in the reduced gel were approximately 30 kDa for E1 and 65 kDa for E2, with the uncleaved E1E2 appearing at approximately 100 kDa. **(B)** Cell count and viability were performed on day 4 post-transfection with the validation genes. Paired t-test shows that there was no significant difference (p > 0.05) in cell viability between the validated genes and the control (empty vector backbone).
